# Whole genome optical mapping reveals multiple fusion events chained by large novel sequences in cancer

**DOI:** 10.1101/166173

**Authors:** Eva K.F. Chan, Desiree C. Petersen, Ruth J. Lyons, Benedetta F. Baldi, Anthony T. Papenfuss, David M. Thomas, Vanessa M. Hayes

## Abstract

Genomic rearrangements are common in cancer, with demonstrated links to disease progression and treatment response. These rearrangements can be complex, resulting in fusions of multiple chromosomal fragments and generation of derivative chromosomes. While methods exist for detecting individual fusions, they are generally unable to reconstruct complex chained events. To overcome these limitations, we adopted a new optical mapping approach, allowing for megabase length DNA to be captured, and in turn rearranged genomes to be visualized without loss of integrity. Whole genome mapping (Bionano Genomics) of a well-studied highly rearranged liposarcoma cell line, resulted in 3,338 assembled haploid genome maps, including 101 fusion maps. These fusion maps represent 175 Mb of highly rearranged genomic regions, illuminating the complex architecture of chained fusions, including content, order, orientation, and size. Spanning the junction of 151 chromosomal translocations, we found a total of 32 Mb of novel interspersed sequences that were not detected from short-read sequencing. We demonstrate that optical mapping provides a powerful new approach for capturing a higher level of complex genomic architecture, creating a scaffold for renewed interpretation of sequencing data of particular relevance to human cancer.

---

Structural variation (SV) is a type of genomic instability characterized by deletions, insertions, inversions, and translocations of large genomic fragments (> 1 kb). Although SV is a form of natural polymorphism (Sudmant et al. 2015; The 1000 Genomes Project Consortium 2015), specific or excessive aberrations have been linked to numerous human diseases (Lupski 1998; Weischenfeldt et al. 2013). In cancer, acquired SVs have been used for molecular sub-classification and shown to be predictive of cancer progression and treatment response (Baca et al. 2013; Papaemmanuil et al. 2016; Holland and Cleveland 2012; Andersson et al. 2016; Hicks et al. 2006). Beyond isolated SVs, large complex genomic rearrangements (CGRs) or chained fusions, defined as the aberrant joining of multiple distant genomic regions (Malhotra et al. 2013), have been implicated in 5%-9% of all cancers (Dzamba et al. 2017). Several models of CGR have been proposed, each thought to be preferential in different cancer types. Chromothripsis, characterized by highly localized shattering and re-ligation (fusion) of tens to hundreds of DNA fragments and an oscillating copy number profile, is, for example, present in 25% of bone cancers (Stephens et al. 2011). Chromoplexy, characterized by closed chains of chromosomal translocations, with little to no copy number alteration, is a key characteristic of prostate cancer (Baca et al. 2013). Breakage-fusion-bridge amplification, characterized by cycles of telomere shortening and fusion, has been implicated in almost every cancer type (McClintock 1941). Despite these distinct models, it is clear that many cancers are probably driven by a combination, and likely intermediate modified forms, of these structural rearrangement processes (Zhang et al. 2013; Garsed et al. 2014).

The complexity of chromosomal abnormalities is typically inferred using various statistical and computational approaches, based on data from karyotyping, microarray-based copy number profiling, and whole-genome paired-end sequencing. A major limitation of these methods is their inability to accurately reconstruct the high levels of amplifications and complex patterns of chained fusions. Reconstruction of a rearranged genome (i.e. to “walk” a derivative chromosome) primarily requires the integration of predicted breakpoints, rearrangement signatures (orientation of sequence read alignment), and copy number profile. However, in the more realistic scenario where derivative chromosomes are formed from multiple processes, rearrangement signatures become confounded and algorithmic assumptions breakdown, making it challenging to accurately reconstruct chromosomal rearrangements (Zhang et al. 2013).

In this study, we demonstrate the utility of a non-sequencing approach, namely, optical (genome) mapping, in capturing somatic chained fusions. We validate the feasibility of using the Irys optical mapping system from Bionano Genomics to capture extensive CGRs in a previously reported well-differentiate liposarcoma cell line 778, derived from an elderly woman with retroperitoneal relapse (Pedeutour et al. 1999). Multi-color fluorescence *in situ* hybridization showed substantial translocations in this cell line, while short-read sequencing of two flow-isolated neochromosome (derivative chromosome) isoforms further confirmed extensive rearrangements with a predominance of intra-chromosomal translocations (Garsed et al. 2014). Using whole-genome mapping, not only did we recapitulate the fusion and copy number profiles of this cell line, importantly, we demonstrate this methodology can identify chained fusion events marked by large intervals undetectable by sequencing alone. We show that optical mapping is an invaluable complement to high-throughput sequencing for reconstructing the architecture of chained fusions commonly found in many cancers.

## Results

### Whole-genome optical mapping and structural variation detection

The Bionano platform is a next generation optical mapping system for generating physical maps (genome maps) of kilobases (kb) to megabases (Mb) in length (Fig. 1). This is achieved by fluorescently tagging and imaging endonuclease motif sites of single-molecules in a massively parallel fashion (Hastie et al. 2013). Imaged molecules are digitized and *de novo* assembled into consensus genome maps, each representing the fingerprint of a large fragment of the sample genome. By comparing sample consensus genome maps against a reference genome, large SVs can be readily identified and visualized. The resolution of this technology is dependent on the frequency of the endonuclease motif in the sample genome. For a human genome, this is between 8 to 11 motifs (labels) per 100 kb using the enzyme Nt.BspQI (GCTCTTCN^). To ensure a high level of specificity when merging and aligning molecules, *de novo* assembly is typically performed on a filtered set of molecules longer than 150 kb in length. Note that, unlike high-throughput sequencing, this method does not require DNA fragmentation, insert size selection or amplification, as it is conversely reliant on intact high molecular weight DNA extraction. While this technology has been available for several years, there had been a prevailing difficulty to accurately predict translocations and inversions. However, a recent breakthrough in bioinformatics development has made possible not only the accurate detection of inversions and translocations, but also the phasing of all detected SVs (Hastie et al. 2017).

**Figure 1.**
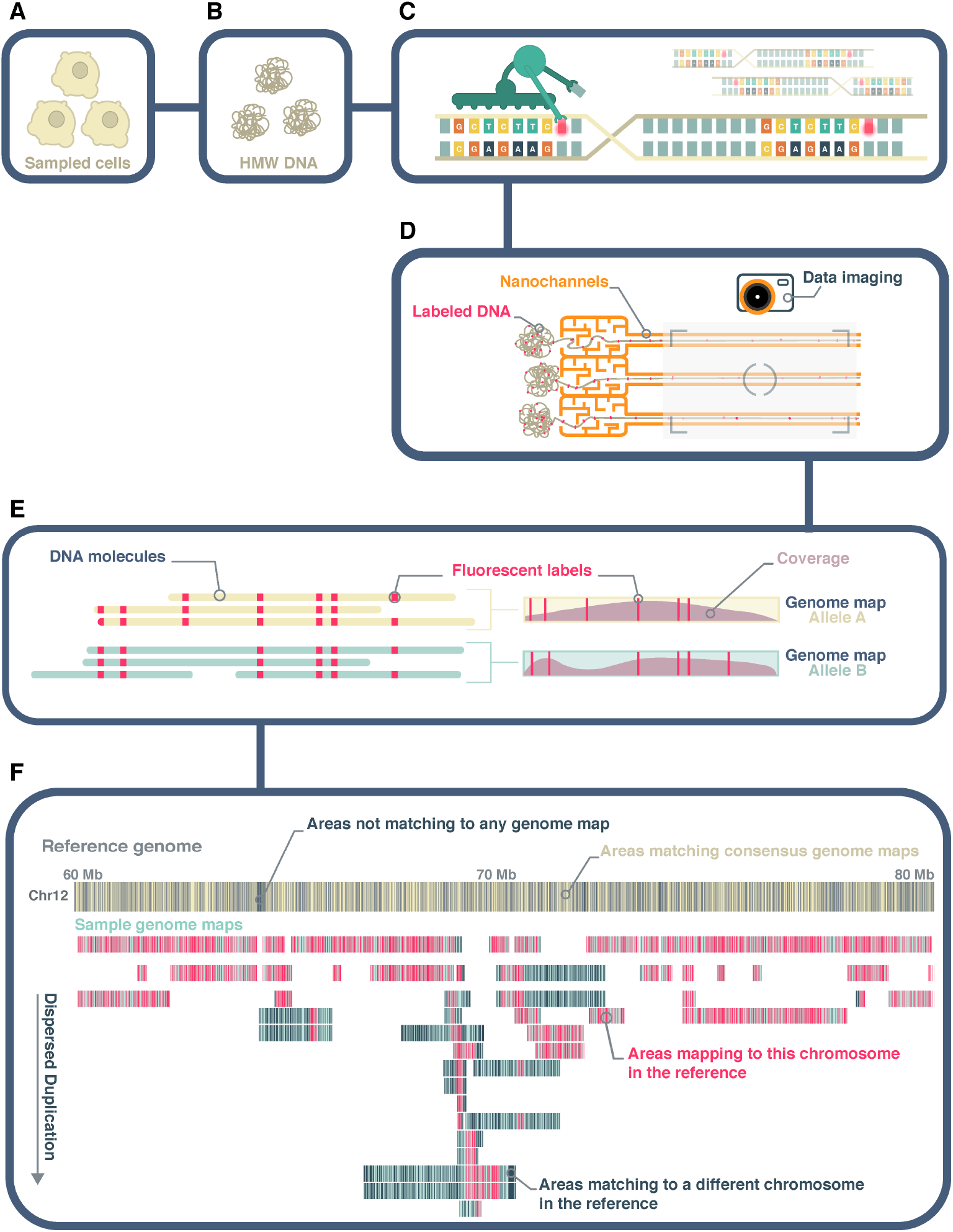
Whole genome optical mapping. (A) & (B) Genome mapping begins with the extraction of intact, high molecular weight (HMW) double-stranded DNA. (C) An endonuclease (Nt.BspQI) that cleaves only one strand of a double-stranded DNA is use to incorporate a fluorescent dye at recognition motifs (GCTCTTCN^) at a density of 8-11 labels per 100 kb. The DNA backbone is also stained with a second fluorescent dye, YOLO-1. (D) Labeled molecules are linearized, flowed into nanochannels, and imaged using the Bionano Irys instrument. (E) Imaged molecules are digitized and bioinformatically assembled into consensus genome maps, and haplotype-phased as appropriate. (F) Structural variations relative to a reference genome are deduced based on differences in size and/or label patterns. Highly rearranged genome maps will align piecewise to multiple reference genomic regions. Conversely, heavily rearranged regions of the reference genome will show alignments from multiple genome maps (dispersed duplication). In this study, the reference genome map is an *in silico* digest of the human reference, hg38. In this figure, sample consensus genome maps are represented as teal colored horizontal bars, overlaid with coverage density plots in mauve, while the hg38 reference is shown as grey horizontal bars. Fluorescent labels of the Nt.BspQI motif are shown as yellow and pink vertical lines overlaid on the reference and sample genome maps respectively. Labels on the reference not aligned to any consensus genome map and labels on sample genome maps not aligned to the displayed reference chromosome(s) are shown in dark blue.

To demonstrate the ability of genome mapping to reveal CGR, we performed whole genome mapping on the 778 liposarcoma cell line(Garsed et al. 2014; Pedeutour et al. 1999). It is important to point out that genome mapping was performed without first isolating the neochromosomes as in the original sequencing study (Garsed et al. 2014). In summary, a total of 798,063 molecules longer than 150 kb were captured and *de novo* assembled into 3,338 consensus genome maps (Table 1; Supplementary File 1). From multi-color fluorescence *in situ* hybridization, we know most chromosomes of this cell line are tetraploid. While the current genome mapping assembly algorithm is agnostic to ploidy greater than two, we were able to deduce an alternate haplomap (haplotype genome map) for 68% (2,271/3,338) of the consensus genome maps. With higher coverage and better assembly algorithm, we can expect more of the consensus maps to be further decomposed into their respective constitutive haplotypes. The average consensus genome map is 1.7 Mb in length with the longest map reaching 6.5 Mb. More than 97% of the genome maps could be aligned to the human reference (hg38) achieving a breadth of coverage of 90%, which is a theoretical limit given the lack of reference genomic information at and around low complexity regions (Fig. 2).

**Table 1.** Genome Mapping Summary. N50: 50% in size of the data is contained within maps or molecules > N50. SV exclude those overlapping N-base gaps and known segmental duplications in hg38, as well as SVs Commonly detected in genome map data across a variety of sample types (Methods).

| Molecule Summary |  |  |
| --- | --- | --- |
| Total number of molecules >150 Kb | 798,063 |  |
| Molecule N50 | 309.5 Kb |  |
| Average label density ( /100 Kb) | 10.3 |  |
| Genome Map Summary | Total | Fusion Maps |
| Number of genome maps | 5,609 | 101 |
| Number of haplomap pairs | 2,271 | 41 |
| Maps without alternate haplotype | 1,067 | 19 |
| Map N50 | 1.377 Mb | 2.377 Mb |
| Map length range | 0.55 - 6.5 Mb | 0.19 - 6.2 Mb |
| Genome maps aligned to hg38 | 97.6% | 100% |
| Genome map length aligned to hg38 | 97.1% | 81.7% |
| hg38 breadth of coverage | 89.9% | 2.1% |
| Structural variation summary | Total | Fusion Maps |
| Deletions (>1 kb, <1 Mb) | 751 | 57 |
| Insertions (>1 kb) | 1,577 | 46 |
| Inversions | 7 | 7 |
| Intra-chromosomal translocations | 34 | 34 |
| Inter-chromosomal translocations | 117 | 117 |

N50: 50% in size of the data is contained within maps or molecules > N50.

**Figure 2.**
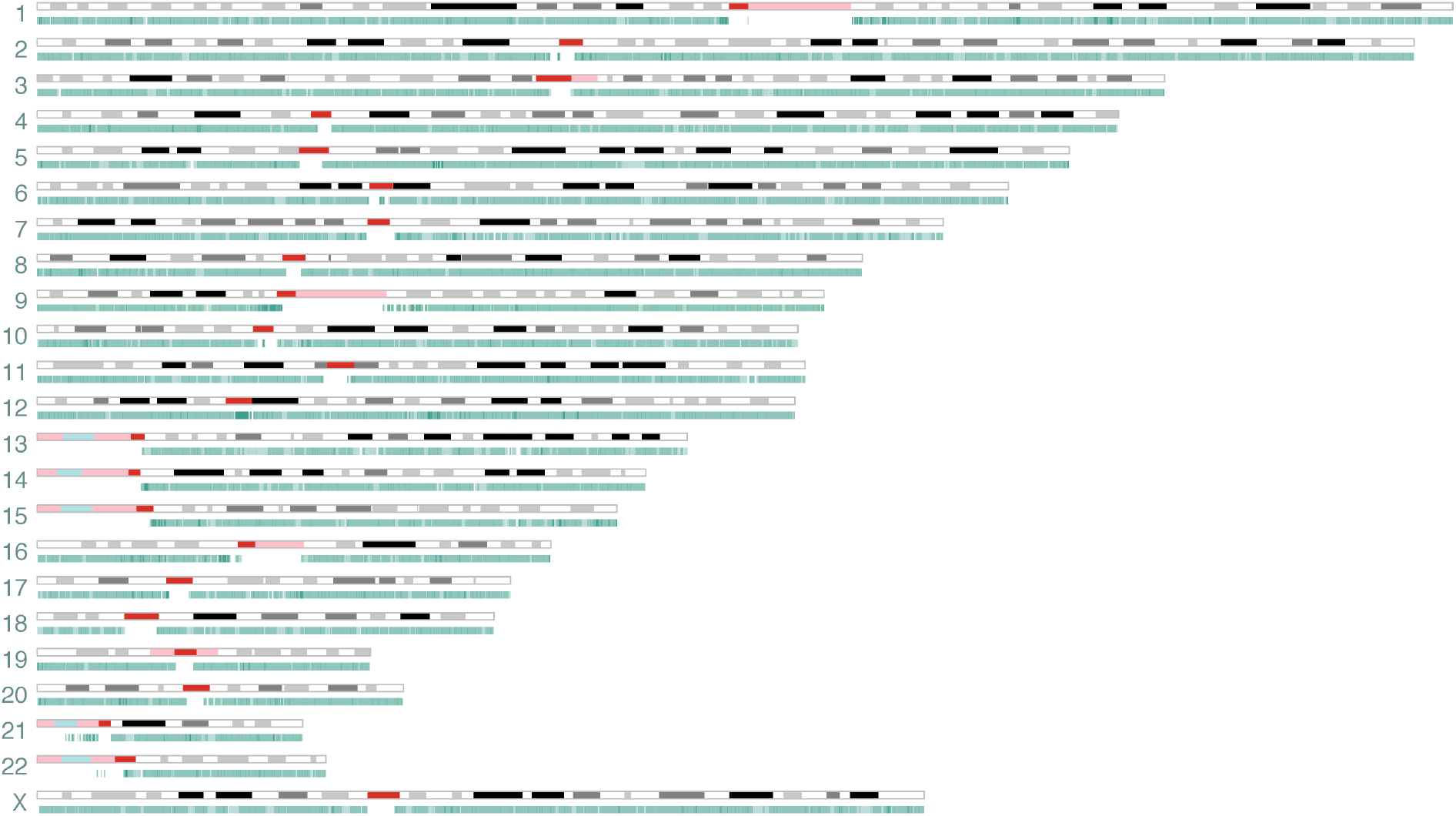
Consensus genome map overview. Shown is an overview of the 3,338 consensus genome maps (teal), from cell line 778, aligned to the human reference genome, hg38. Overlapping alignments to hg38 (i.e. Alignment density) is reflected in the color gradient of the genome maps. The reference genome is represented by its cytogenetic G banding (UCSC Table *cytoBandIdeo*, last updated 11 June 2014) where Dark to Light gray in this figure correspond to Giemsa stain intensity, acrocentric (acen) bands are represented in red, variable heterochromatic region (gvar) in pink, and stalks (tightly constricted regions) in blue. Breadth Of coverage is Theoretically limited to 90% of the reference genome due to uninformative regions, including acen, gva, and stalks.

A total of 2,486 SVs (> 1 kb) relative to hg38 were identified (Table 1). This excluded all SV overlapping N-base gaps and known segmental duplications in hg38, as well as common false positive translocation breakpoints (breakpoints frequently found in genome maps not known to contain translocations; Methods). We observe twice as many large insertions (1,577) compared to deletions (751). This is similar to previous reports for other cancer cell lines using the same technology, reporting between 1.5 and 2.7 fold more insertions than deletions(Dixon et al. 2017; Zook et al. 2016) (Fig. 3A) and for non-cancer samples (data not shown). Although insertions are more abundant, they are generally smaller in size (median: 2.7 kb, interquartile range (IQR): 1.7-4.9 kb) relative to deletions (median: 3.5 kb, IQR: 2.2-6.3 kb) (Fig. 3B). The largest insertion observed is 386 kb within Chr1: 172,422-172,618 kb, which upon inspection is, in fact, a replacement of the 196 kb interval with a chained fusion of 84.2 kb fragment from chromosome 1 (188.314-188.394 Mb), a 143.6 kb fragment from chromosome 15 (98.430-98.573 Mb) and a 354 kb fragment that does not align to hg38 (see below for further discussion on unaligned genome map fragments) (Supplementary Fig. S1A). The largest deletion of 850 kb at Chr15: 90,881-91,870 kb was detected in two pairs of haplomaps, though we note genome maps spanning the deleted interval are also present (Supplementary Fig. S1B), suggesting this genomic fragment is not lost to the cancer genome. The distribution of large insertions and deletions is roughly random across the genome, with the proportion of each SV type being linearly correlated to chromosome size (Fig. 3C).

**Figure 3.**
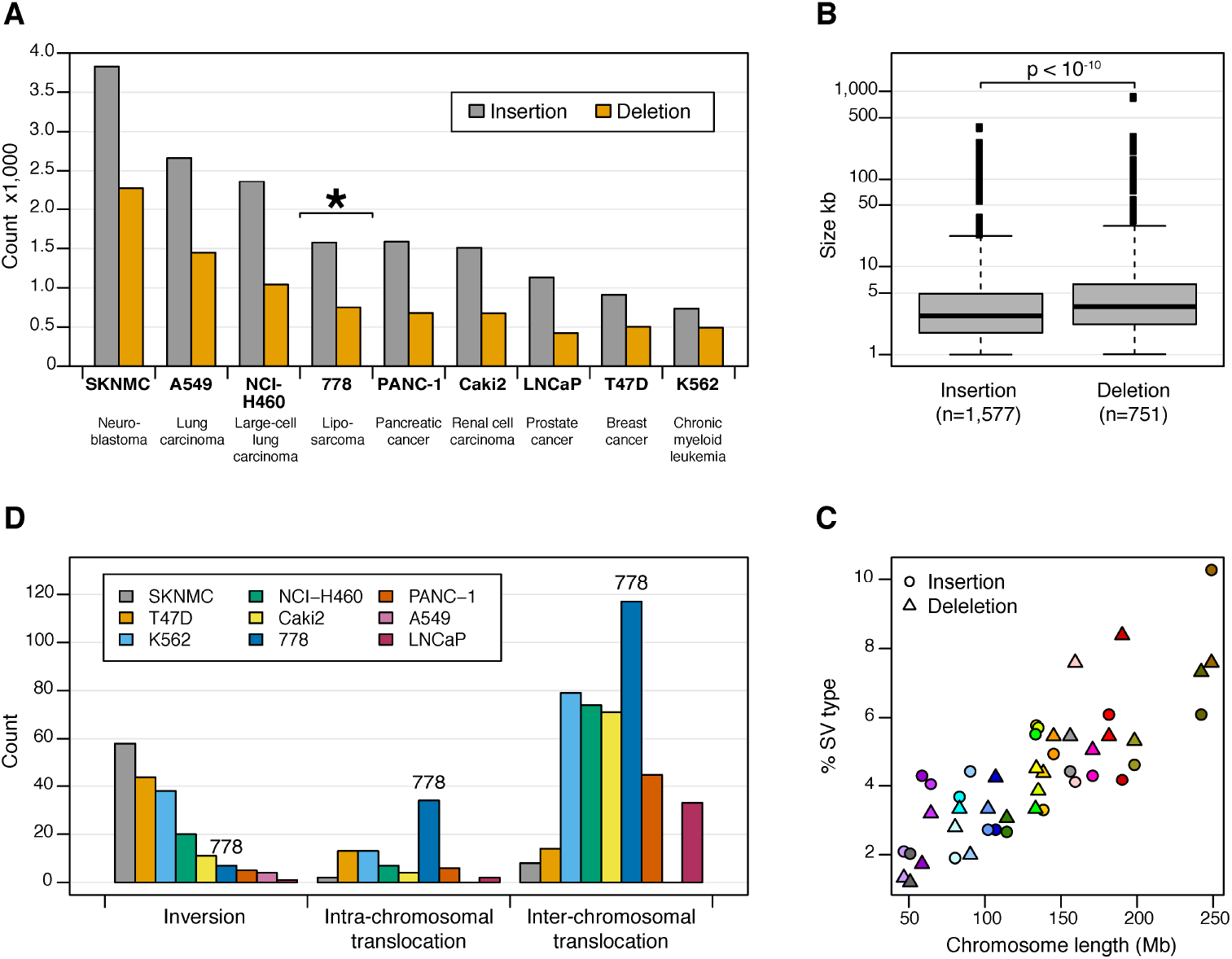
Structural variations Identified using whole-Genome optical mapping. (A) & (D) Comparison Of large structural variations identified in liposarcoma cell line 778 with eight other cancer cell lines reported in Dixon *et al.* 2017. All SVs from both studies were determined using the Bionano Irys optical mapping system. Panel (A) shows the typical 1.5 To 2.7 fold More insertions Relative to deletions, while panel (D) shows cell line 778 harbor much more intra-and inter-chromosomal translocations relative to other cancer cell lines. (B) Boxplot showing a statistically significant difference between deletion and insertion sizes, for SVs larger than 1 kb. Thick black horizontal lines in the middle of the boxplots correspond to median values, whileshadedgray boxes encompass the interquartile ranges. (C) The Proportions of Large insertions and deletions found in cell line 778 are correlated with chromosome length.

Besides large deletions and insertions, we identify 151 translocations (34 intra-and 117 inter-chromosomal) and five inversions across seven genome maps (Fig. 3D). As observed for the large deletions and insertions, the total number of inversions most closely represents that found in the renal and pancreatic cell lines using the Bionano Irys technology (Dixon et al. 2017). The total number of inversions observed within a single cancer cell line ranges from one in the prostate cancer cell line LNCaP to 68 in the neuroblastoma cell line SKNMC. In contrast, we observe considerably more chromosomal translocations in 778 than has previously been reported for other cancer cell lines. Dixon *et al.* 2017 reported, at most, 92 translocations (13 intra-and 79 inter-chromosomal) in the chronic myeloid leukaemia cell line K562, while no translocation was found in the lung carcinoma cell line A549. This latter report for A549 is surprising as extensive CGRs have previously been shown (Peng et al. 2010), and the lack of translocation in this recent report may be explained by differences in sub-line passages of this relatively old cell line (Gu et al. 2016). Thus, optical mapping is capable of capturing a rich diversity of translocations in carcinogenesis.

### Fusion maps and chained fusions

To better interrogate CGR, we focused on 101 genome maps, which included all observed translocations and inversions, as well as insertions and deletions whose breakpoint pairs on the reference genome are more than 100 kb apart (Methods). This set of genome maps, termed “fusion maps”, includes 41 haplomap pairs and 19 singletons, totaling 175.5 Mb diploid length. Supplementary Figure S2 displays the schematic representation of the full list of 101 fusion maps, from which three examples are shown in Figure 4. Alignment of the fusion maps to hg38 revealed a total of 282 SVs, with the majority being inter-chromosomal translocations, followed by large deletions, insertions, intra-chromosomal translocations and inversions (Table 1). Half of the fusion maps have at least one inter-chromosomal translocation and one quarter have two or more (Supplementary Fig. S3). In contrast, although 47% of the fusion maps contain at least one deletion, the majority has no more than one. Surprisingly, the typical 1.5 to 2.5 ratio of insertions to deletions was not observed in the fusion maps, which contain more deletions (69) than insertions (55), though deletion sizes remain significantly larger than insertions (Supplementary Fig. S4). This new observation alludes to different mechanisms and selective pressures driving the high frequency of insertions in many cancer and non-cancer samples relative to CGR-type insertions.

**Figure 4.**
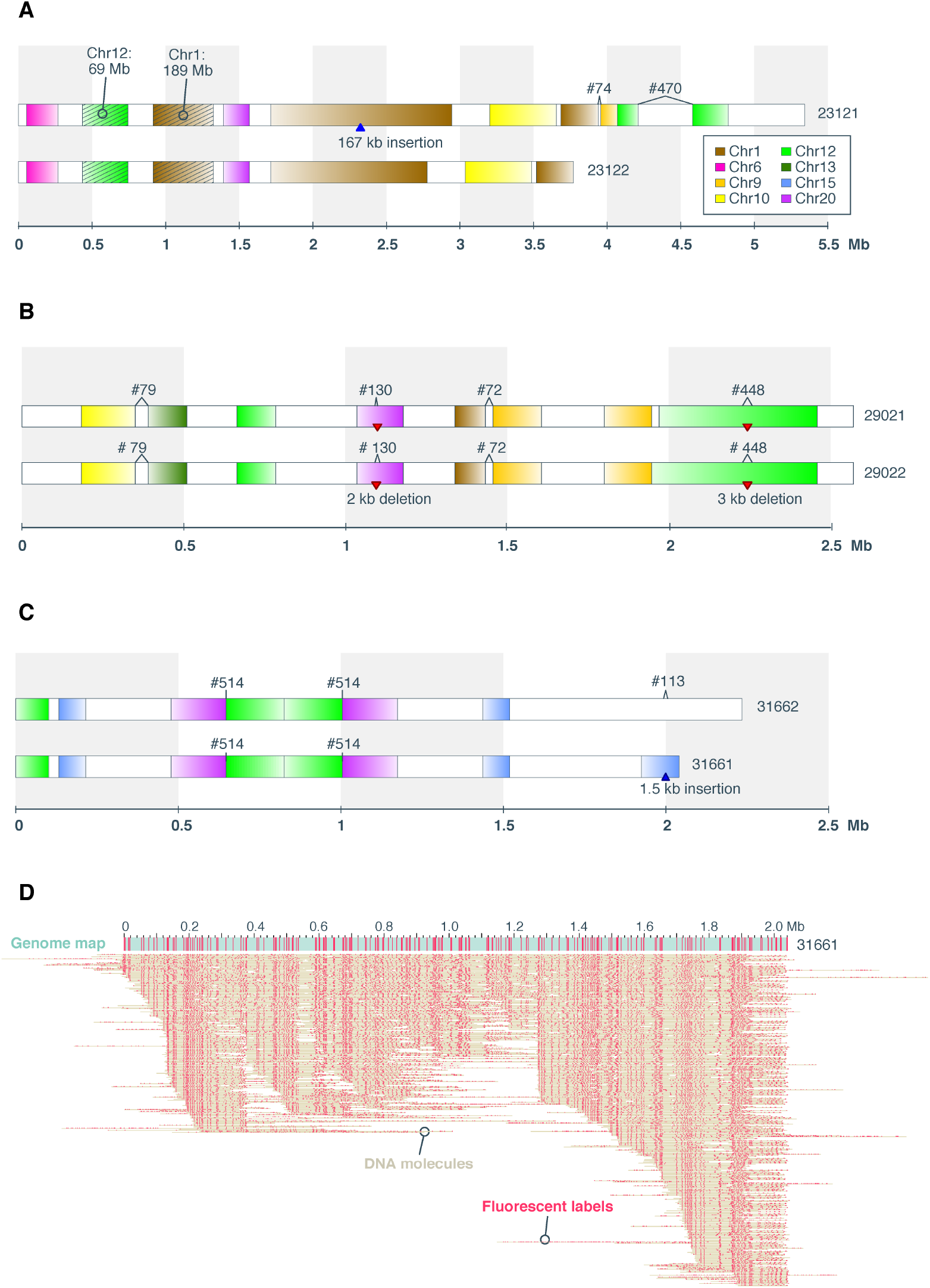
Fusion map examples. Panels (A), (B), and (C) Are schematics Of three fusion Haplomap pairs Containing complex Genomic rearrangements. Genome map sizes are indicated on the Horizontal axis, in megabase units. In each panel, Fragments aligning to hg38 chromosomes Are indicated by the default UCSC Chromosome color Scheme (color key in panel (A)). Uncolored (white) intervals Correspond to regions not aligned to the reference. Alignment orientation to hg38 is indicated by Dark to light color gradient corresponding to 5’ to 3’ alignment to the Positive strand. Deletions and insertions are Indicated by red downward triangles and blue upward triangles respectively. The two most frequently represented reference fragments (Chr1: 188,188,529-189,139,998 and Chr12: 68,713,897-69,940,974) found in the fusion maps are shown with diagonal strips and indicated as Chr1:189 Mb and Chr12:69 Mb. Previously identified fusions from sequencing data are numbered per Supplementary Table S1 (c.f. Garsed et al.2014) and indicated above the genome maps. Panel (D) shows the molecules Aligning to, and making up, Consensus genome map #31661, which contains an inverted chained fusion as shown in panel (C). Here, The 2 Mb consensus genome map is represented by a teal horizontal bar following the convention in Figure 1. Individual molecules are represented as “dots on a string”, where each yellow horizontal line represents a molecule and pink dots represent fluorescent labels. Panels (A) to (C) are A subset of the 101 Fusion maps shown In Supplementary Figure S2.

Further interrogation of the fusion maps revealed extensive chained fusions (Supplementary Fig. S2). Highly duplicated and transposed reference genomic regions are notable from the pileup of genome maps aligning to these regions (Fig. 1). Chromosomes 12 and 1 are most represented, contributing 21% and 15% of all reference genome fragments found in the fusion maps, respectively (Fig. 5). Specifically, regions Chr12: 68,713,897-69,940,974 and Chr1:188,188,529-189,139,998 were found in more than 10 fusion map pairs (Methods; see example in Fig. 4A). In contrast, contributions from four chromosomes 2, 11, 18, and 22 are notably absent. Almost all chromosomes represented in the fusion maps are impacted by translocations (Fig. 5). Of the 117 inter-chromosomal translocations, 58% involve chromosomes 1, 12, and 15. Specifically, breakpoints of these translocations are largely localized to two loci on chromosome 1: 185.7-186.7 Mb and 188.7-189.7 Mb), Chr12: 68.7-70.9 Mb, and two loci on Chr15: 91.8-92.9 Mb and 98.2-98.6 Mb). For intra-chromosomal translocations, nearly 79% involve chromosomes 12, 1, and 7. While breakpoints involving chromosomes 12 and 1 are predominantly localized (Chr12: 69 Mb, Chr12: 92 Mb, Chr1: 172 Mb, Chr1: 189 Mb), breakpoints on chromosome 7 are more diffused (Supplementary Fig. S5). Of note are the highly represented reference donor loci, Chr12: 69 Mb and Chr1: 189 Mb, which also appear to be hotspots for both intra-and inter-chromosomal translocations.

**Figure 5.**
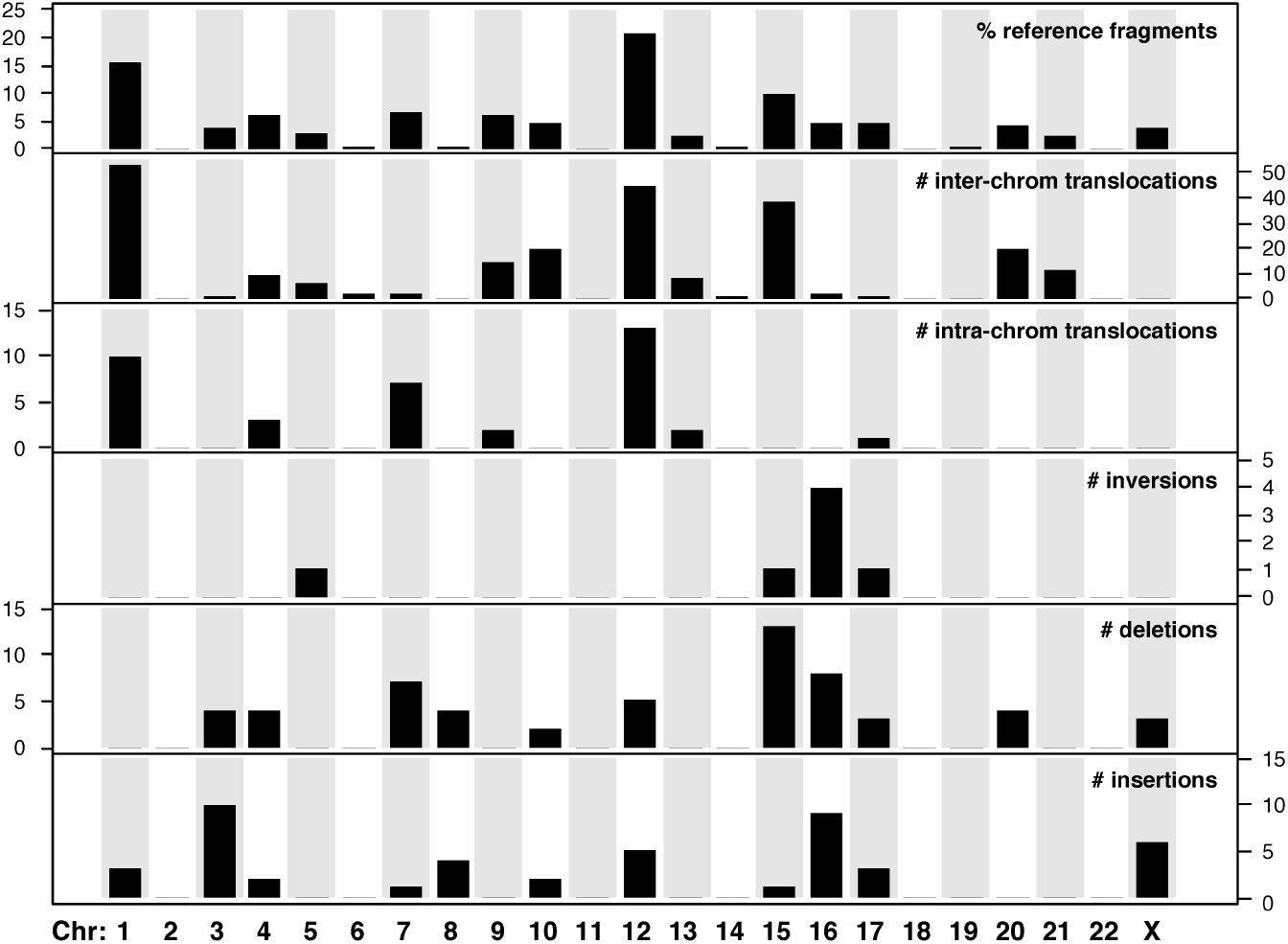
Chromosomal distribution of donor sequences and five structural variation types observed in the 101 fusion maps. Top panel shows the percent of reference donor fragments found in the fusion maps belonging to each of the 22 autosomes and chromosome X. The next five panels show the numbers of each SV event involving the corresponding chromosomes.

An inversion is typically signified by two breakpoints in the reference genome and two fusion junctions in the rearranged genome map (Fig. 6A). In three of the inversions, however, only one fusion junction was captured, thus, we can only speculate on their size (Supplementary Fig S6). In contrast, both fusion junctions of an inversion at Chr16: 70,195,951 -74,372,427 are captured by two haplomap pairs, allowing the 4 Mb inversion to be fully defined (Fig. 6B). The remaining inversion is found on a genome map containing an inverted duplication of a series of fusion events (map #31661; Fig. 4C), further discussed below.

**Figure 6.**
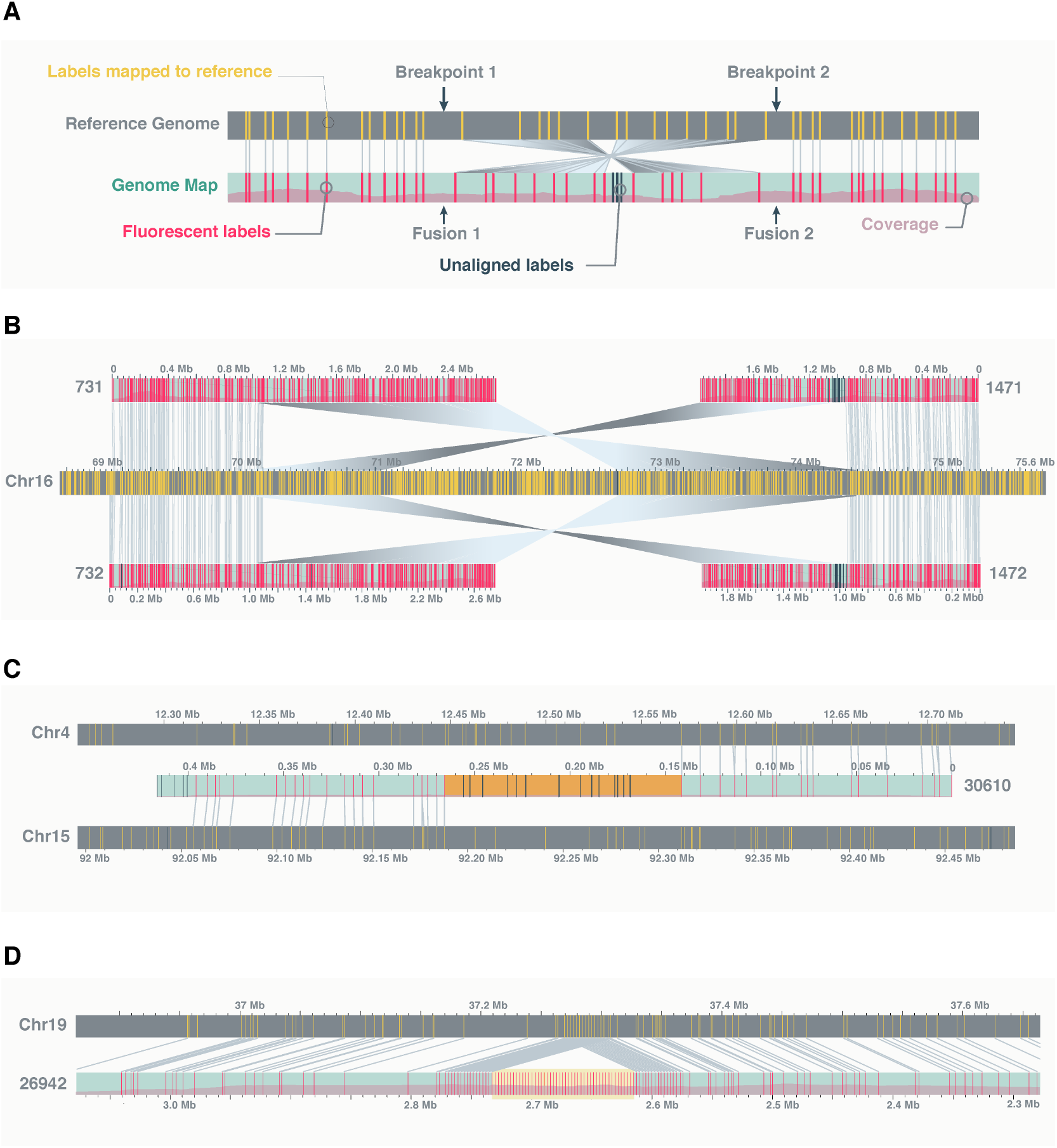
Structural variation examples. (A) A genomic inversion is ideally characterized by two breakpoints in the reference genome and two fusion junctions in the rearranged genome. (B) An Example of a fully resolve 4 Mb inversion, characterized by two pairs of optical maps, each carrying one fusion junction with flanking fragments aligning in opposing direction to one side of the two reference breakpoints. (C) An example of a translocation between chromosomes 4 and 15, showing “complex” label patterns at the rearrangement junction (highlighted in orange). (D) An example of a 159 kb insertion, showing “repeat” label patterns of the inserted fragment (highlighted in yellow). In this case, the repeat corresponds to the SST1 satellite. Additional examples Highlighting complex labels patterns at translocation junctions relative to repetitive label patterns of insertions can be found in Supplementary Figure S7. The schema In this figure follows the same convention as outlined in Figure 1. Matching labels Between sample And reference Genome maps are Connected by grey lines. For clarity, any additional genome maps aligning to the reference regions of interest are hidden from view.

Most deletions (23%) on the fusion maps involve chromosome 15 followed by chromosomes 16 (14%) and 7 (12%) (Fig. 5). For large insertions, the majority involves chromosomes 3 (22%), 16 (20%), and X (13%). Aside from a substantially reduced incidence of large insertions and deletions on chromosomes 1 and 12, the distributions of these two SV types largely mirror those for intra-chromosomal translocations. Together these observations can be attributed to the rearrangement processes underlying the evolution of this cancer cell line, which included an initial catastrophic event causing the fusion of specific genomic fragments forming the cores the derivative chromosomes (see next section below).

A key observation from this study is how clearly chained fusions can be visualized. Features such as order, orientation, and size of rearranged DNA fragments are readily observed (Supplemental Fig. S2). More importantly, and unprecedented from sequencing data, patterns and sizes of fusion junction intervals are also captured. These intervals represent unknown fragments that do not align to the reference genome, and total approximately 32 Mb across the 101 fusion maps. These unmapped fragments likely correspond to regions of significant rearrangements rather than repetitive elements, as their label patterns are typically “unique” (examples in Fig. 6C and Supplementary Fig. S7A). This is in contrast to insertions that tend to involve repeat elements (Fig. 6D; Supplementary Fig. S7B). Other features identifiable from optical mapping include large inverted duplication, which can be difficult to detect from sequencing data. Surpassing this complexity, we identified an inverted duplication of a chained fusion t(15;20;12) (Fig. 4C), suggesting the duplication occurred after the chained fusion. Manual examination of the molecules making up the consensus genome map confirmed assembly artifact is unlikely (Fig. 4D).

### Fusion maps and neochromosomes

There are several lines of evidence suggesting that, although genome mapping was performed on the entire 778 cell line, the 101 fusion maps identified largely correspond to the two neochromosome isoforms previously sequenced (Garsed et al. 2014). Firstly, CGR and copy number profiles of the fusion maps strongly recapitulate those derived from the two neochromosome isoforms (Garsed et al. 2014) (Fig. 7). A total of 26 (4.4%) of the 587 previously reported neochromosome fusions are also present in 35% of fusion maps, while 54 (18.4%) of the 282 CGRs identified in the 101 fusion maps were also identified from short-read sequencing of the neochromosomes (Supplementary Table S1). For the mapped fusions, the majority (39%) of concordances are intra-chromosomal translocations follow by inter-chromosomal translocations (21%) and deletions (19%). An obvious reason for the low overlap rate is related to our definition of fusion maps, which excluded maps without relatively large CGR (reference breakpoint distances > 100 kb) or maps with only SV overlapping N-base gaps or known segmental duplications (see above and Methods). Of the previously reported neochromosome fusions, 71 (12%) have breakpoint distances <50 kb and 81 (14%) overlapped with segmental duplications.

**Figure 7.**
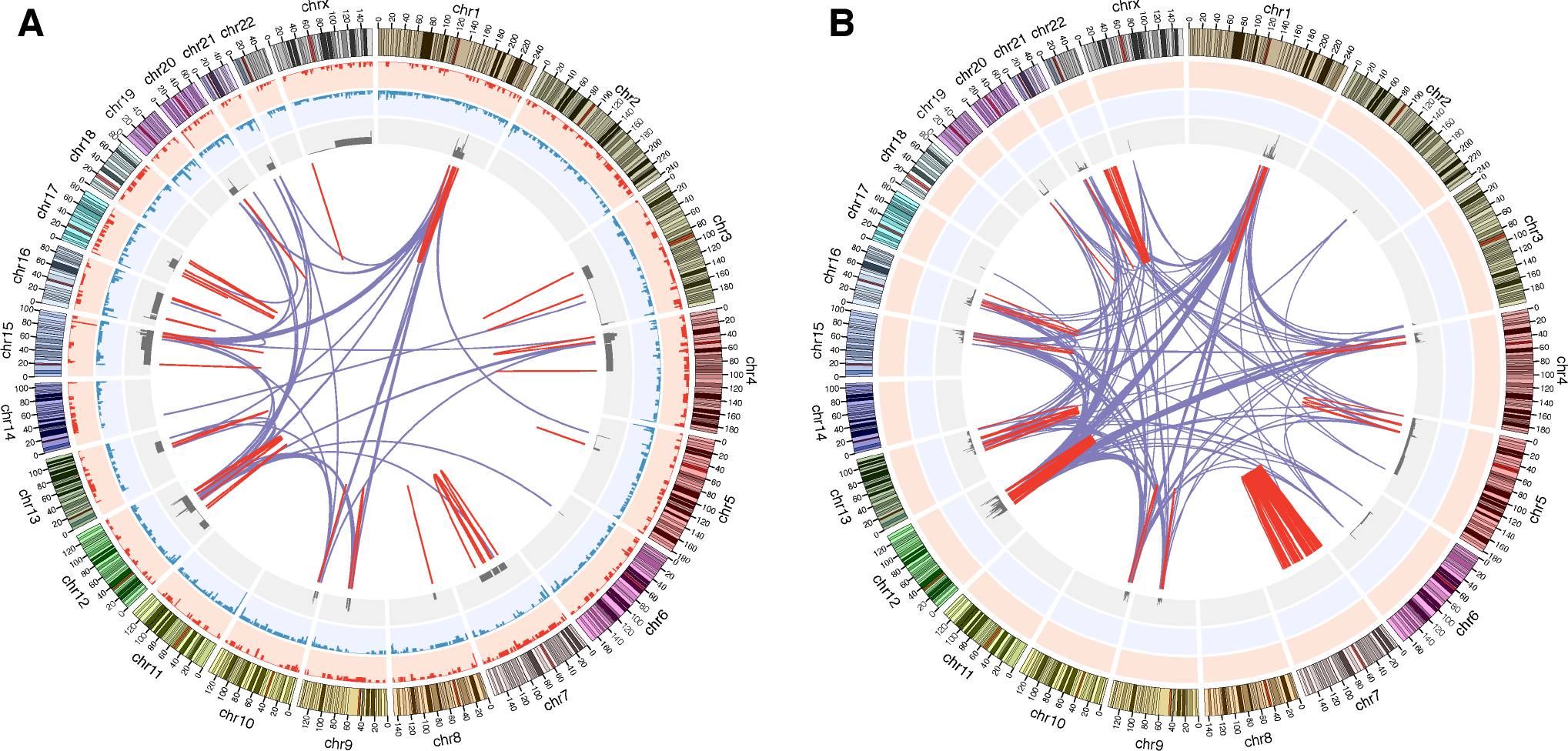
Circos plots of genomic variations in cell line 778. (A) Circos plot derived from whole genome mapping of the cell line. (B) Circos plot derived from short-read sequencing of two neochromosome isoforms, using data from Garsed et al.15. For both plots, moving inward from the outer ideogram are histograms of deletions (>1 kb; red) and insertions (>1 kb; blue) in the background genome. For the neochromosome circos plot (B), these two tracks are only placeholders, as the background genomes were not sequenced in the original study. The next track, in gray, shows copy number profiles of the fusion maps (A) and neochromosomes (B). Linked lines in the middle of the circos plots show complex genomic rearrangements: red for intra-chromosomal and purple for inter-chromosomal translocations.

A second line of evidence suggesting the fusion maps may represent much of the 778 neochromosomes is in the absence of DNA fragments from chromosomes 2, 11, 18, or 22 in the fusion maps (Fig. 5). This is largely consistent with the absence of both genomic fusions and contiguous genomic regions (defined as contiguous and interconnected fragments of the reference genome present in the neochromosomes) involving chromosomes 8, 11, 14, 18, and 19 in the neochromosomes (Garsed et al. 2014). The absence of chromosome 22 in the fusion maps is likely because this chromosome is relatively un-rearranged with the exception of a small cluster of intra-chromosomal rearrangements found in one of the neochromosome isoforms, suggesting fusions involving chromosome 22 are late events acquired after the formation of the neochromosome cores. Of the three chromosomes (8, 14, 19) present in the fusion maps but absent in the neochromosomes, we note, in each case, only one fusion map (or map pair) contains the corresponding chromosome fragments. For the single haplomap pair (maps #24711/2; Supplemental Fig. S2) containing a chromosome 8 fragment, while a 155 kb deletion is present, no translocation was identified. Similarly, only one fusion map (#23051) aligned to chromosome 14; a fusion map composed of a 160 kb chromosome 3 fragment fused to a 114 kb fragment from chromosome 14. For these two cases, the corresponding fusion maps may be representative of the background genome, not the neochromosomes. For the last case, again, only a single fusion map (map #27361) was aligned to chromosome 19. This 2.9 Mb fusion map is primarily composed of the telomere of the long arm of chromosome 3, which appears to share 56 kb of label similarity with the telomere of the short arm of chromosome 19. Therefore, the inclusion of this chromosome 19 fragment in the fusion map is most likely an alignment artifact (Supplementary Fig. S8).

A third line of evidence is in the dense clustering of genomic rearrangement breakpoints previously shown in the sequencing data and apparent in the fusion maps (Fig. 8). Such clustering of fusion breakpoints is characteristic of chromothripsis (Korbel and Campbell 2013). Specifically, it was theorized that the 778 neochromosomes likely resulted from an initial chromothriptic even involving chromosome 12 leading to (i) the majority of chromosome 12 contiguous genomic regions being intra-chromosomal, (ii) chromosome 12 being the dominant inter-chromosomal translocation partner, and (iii) the subsequent amplification of various chromosome 12 genes by the breakage-fusion-bridge mechanism. Consistent with this, we note in the fusion maps, that (i) 38.2% of all intra-chromosomal translocations and (ii) 38.5% of all inter-chromosomal translocations involve chromosome 12, and (iii) one of the two most represented reference donor fragments in the fusion maps is Chr12: 68-70 Mb, encompassing genes *NUP107*, *YEATS4*, *MDM2*, *SLC35E3*, and *BEST3*.

**Figure 8.**
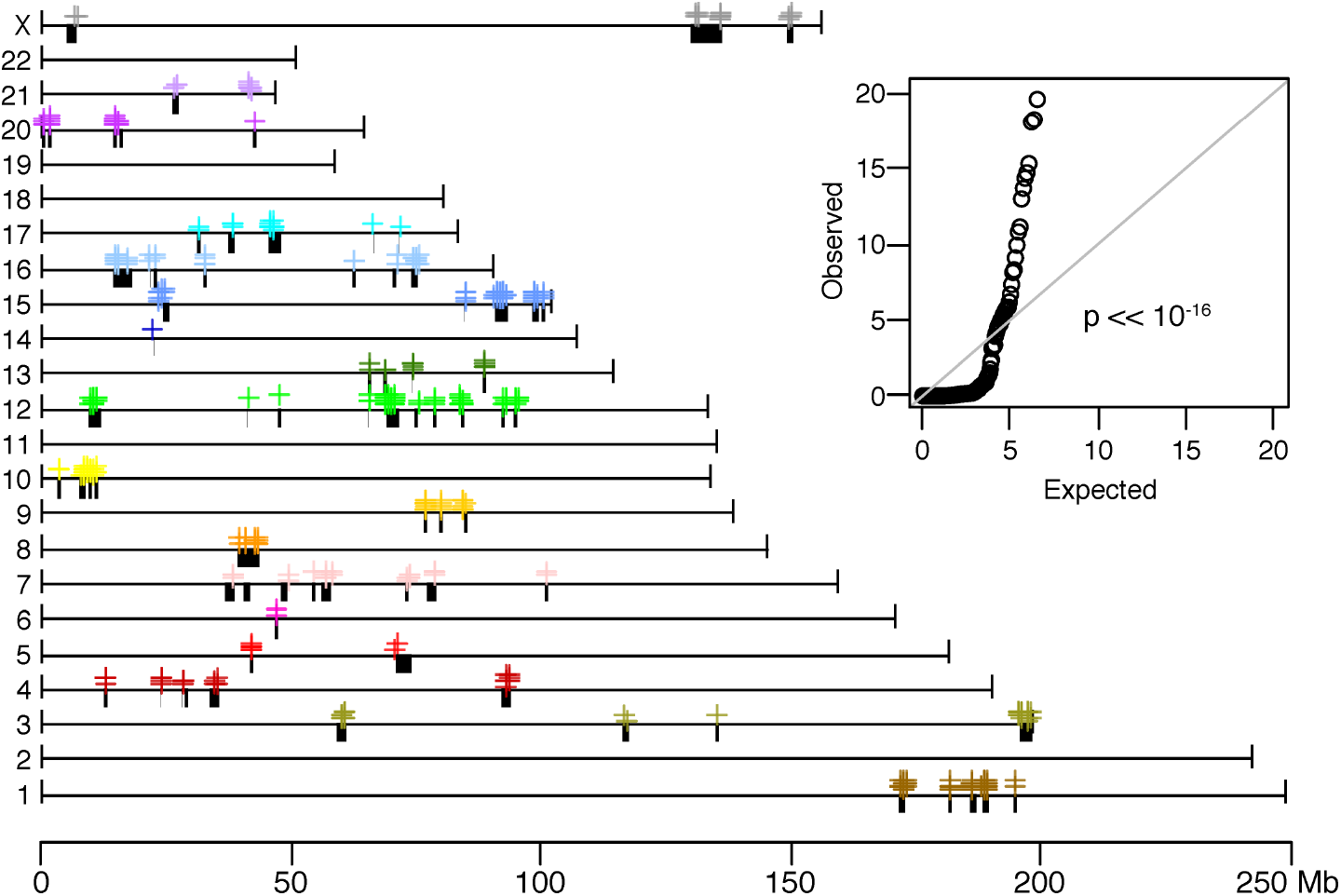
Fusion clusters. Donor fragments (black bars) and complex genomic rearrangement breakpoints (colored crosses) found in the 101 fusion maps are highly localized on the reference genome map, hg38. Fusion breakpoints on some chromosomes are notably at the boundaries of reference donor fragments (e.g. chromosomes 5 and X), suggesting the acquisitions of these fragments in the rearranged genomes were late events. This is in contrast to donor fragments with “internal” CGRs (e.g. chromosome 12) suggesting their acquisitions were early events. The inset figure is a quantile-quantile plot of the observed adjacent fusion breakpoint distances relative to the null expectation of random distribution, indicating the distribution of fusion breakpoints is statistically significantly non-random. Colors of plotting symbol correspond to the default UCSC chromosome color scheme.

Based on overlap analysis, we identified 100 reference donor fragments from the 101 fusion maps totaling 64 Mb (Methods). In contrast, the two 778 neochromosome isoforms is made up of 23 Mb of the reference donor genome (Garsed et al. 2014). Thus, it is plausible that some of the fusion maps identified represent the background (albeit rearranged) genome, which is consistent with observed chromosomal translocations in 778. Finally, we note that genome map fusions not previously identified from short-read sequencing tend to have significantly larger intervals between fusion points on the rearranged genome (average 112 kb with 64% > 20 kb; Supplementary Fig. S9). This is exemplified in Figure 4B, where four fusion events, including two intermediate sized deletions (2 - 3 kb) were previously identified from sequencing. Our inability to detect these large rearrangement junctions can primarily be explained by two compounding issues. One is the high level of genomic amplification followed by rearrangement that commonly occurs in cancer, and which causes breakpoints to be present in different, potentially non-adjacent, copies. This is exacerbated by the need to fragment the genome to “manageable” sizes, typically much less than 20 kb, prior to short-read sequencing (Korbel and Campbell 2013; Guan and Sung 2016). Together, this effectively prevents large fusion intervals to be captured and resolved by short-read sequencing. While bioinformatics methods, such as *de novo* breakpoint assembly, may overcome this technology-specific bias, these approaches are difficult with high false negative rates (Malhotra et al. 2013). In contrast in whole genome mapping there is no deliberate shearing or fragmentation step and meticulous care is taken to ensure DNA molecules of hundreds of kilobases to megabases in length are captured.

## Discussion

In this study, we demonstrated the utility of whole genome mapping to reconstruct unresolved complex architectures of chained fusions. High throughput sequencing has highlighted the importance of CGR in many human cancers. While sequencing can identify simple SVs down to single-base variant resolution, determining the full spectrum of large CGRs remain difficult. In particular, sequencing cannot accurately reveal the physical relationships of fusion events. This is largely because CGRs are composed of multiple, often overlapping, SVs that tend to result in confounding rearrangement signatures (Zhang et al. 2013; Liu et al. 2015). This challenge is further exacerbated by inter-cell heterogeneity and aneuploidy, both hallmarks of cancer genomes. Optical mapping is designed to capture single molecules of hundreds to thousands of kilobases in size, allowing large rearrangements to be observed and phased (Jaratlerdsiri et al. 2017; Dixon et al. 2017; Hastie et al. 2017). As such, we speculate this method can provide an ideal complement to genome sequencing for resolving complex genomic architectures. We elected to test our hypothesis on a well-described, highly rearranged liposarcoma cell line 778, with known derivative chromosomes.

We performed whole genome mapping of 778, generating 3,338 assembled haploid genome maps of up to 6.5 Mb in length, and recovering 101 fusion maps representing the most highly rearranged regions of the cancer genome. For the first time, chained fusions can be directly observed without the need for complex algorithmic inferences reliant on intricate assumptions. Using this method, 64% of the fusion events were found to have rearrangement junctions larger than the typical short-read sequencing fragmentation length of 20 kb. Furthermore, we uncovered a total of 32 Mb genomic fragments (ranging from 40 bp to 2.6 Mb) within fusion junctions that are apparently distinct from the reference genome. We believe these unaligned regions represent highly degenerate genomic fragments. The impact and function, if any, of these novel regions remain to be explored.

We previously proposed a process of liposarcoma neochromosome formation, initiated by chromothripsis of chromosome 12 leading to the formation of a circular chromosome, followed by a series of breakage-fusion-bridge cycles and punctuated chromothriptic events, and ultimately terminating in a stabilized linear form (Garsed et al. 2014). The non-random distribution of translocations observed in this study concurs with the assumption of a single catastrophic chromothriptic like event, occurring early in the evolution of the 778 cancer cell line. At the same time, the involvement of almost all chromosomes implies additional, plausibly stepwise, rearrangement processes along with post-hoc selection were also at play following the initial catastrophic genomic event. Combining optical mapping with previous information from short-read sequencing and karyotyping, we have gained further insight into the formation and genomic architecture of this highly rearranged cancer cell line.

Complex genomic rearrangements has major roles in cancer development, emerging as a major indicator of cancer subtype, progression and treatment response (Baca et al. 2013; Papaemmanuil et al. 2016; Holland and Cleveland 2012; Andersson et al. 2016; Hicks et al. 2006). Comprehensively detecting CGR in cancer remains challenging. No single approach can comprehensively identify all SV, as each approach has their strengths and weaknesses. Integrative methods combining different data types are showing promise (Dixon et al. 2017; Mostovoy et al. 2016). As long-range sequencing remains cost prohibitive, optical mapping provides a logical intermediate between traditional low-resolution karyotyping and current short-read sequencing.

## Methods

### Cell culture

Approximately 10 million cells of the lineage 778 were provided by A. Cipponi (Garvan Institute). This cell line was derived from a retroperitoneal relapse of a well-differentiated liposarcoma from a 68-year-old woman in 1993 (Pedeutour et al. 1999).

### Optical mapping

High molecular weight DNA was isolation using IrysPrep Plug Lysis Long DNA Isolation Protocol (Bionano Genomics). In brief, cells were trypsinized, washed in FBS/PBS, counted, rewashed in PBS and embedded in agarose plugs using components from the Bio-Rad Plug Lysis Kit. The plugs were subjected to Proteinase K digestion (2 x 2h at RT). After a series of washes in buffer from the Bio-Rad kit, followed by washes in TE (Tris-EDTA), the plugs were melted and treated with GELase enzyme (Epicenter), The high molecular weight DNA was released and subjected to drop dialysis. The DNA was left to equilibrate for four days, then quantification using the Qubit Broad Range dsDNA Assay Kit (Thermo Fisher Scientific).

Using the IrysPrep NLRS assay (Bionano Genomics), 200-300 ng/μL of high molecular weight DNA underwent single-strand nicking with 10U of Nt.BspQ1 nickase (New England BioLabs). Nicked sites were repaired with fluorophore-labeled nucleotide to restore strand integrity. The backbone of fluorescently labeled double-stranded DNA was stained with the intercalation dye YOYO-1. Labeled molecules were directly loaded onto IrysChip^®^, without further fragmentation or amplification, and imaged using the Irys^®^ instrument.

Imaged molecules were digitized, and raw molecules filtered based on minimum molecule length of 150 kb and a signal-to-noise ratio between label and background fluorescence greater 2.75. Further filtering and de novo assembly of single-molecules into consensus genome maps were performed with the Bionano Solve™ v3.0 software (Lam et al. 2012; Hastie et al. 2013, 2017). An advantage of the new software is that it is “haplotype-aware”, with the ability to reconstruct heterozygous haplomaps.

### Structural variation detection

Structural variations were identified relative to the human reference genome, hg38, whose genome maps were bioinformatically deduced based on predicted Nt.BspQI motif sites. Structural variation detection was performed using the Bionano^®^ custom SV caller (https://bionanogenomics.com/wp-content/uploads/2017/03/30110-Rev-B-Bionano-Solve-Theory-of-Operation-Structural-Variant-Calling.pdf). In brief, consensus genome maps were aligned to hg38 and regions with mismatch label patterns, such as loss/gain of labels and changes in inter-label sizes, were identified as corresponding to potential structural variations.

All SVs spanning known N-based gaps or segmental duplications (per the UCSC *gap* (last updated: 24 Dec 2013) and *genomicSuperDups* (last updated 14 Oct 2014) tables) in hg38 were excluded. Further, translocations spanning any of the 142 “common translocation breakpoints” were excluded. This list of common breakpoints was provided by Bionano and includes translocations observed in at least four of 147 genome map assemblies generated from normal control human samples. Because of the resolution of the Bionano optical mapping system, insertions and deletions smaller than 1 kb were not included in this study. The size of an SV is taken as the absolute difference between the SV intervals on the sample and reference genome maps.

The histograms of insertions and deletions in the Circos plots of Figure 7 were calculated as the total number of insertions and deletions within 1 Mb bins.

### Fusion genome maps

Fusion maps were defined as consensus genome maps with at least one complex genomic rearrangement, including all translocations and inversions as well as insertions and deletions whose hg38 reference breakpoint pairs are farther than 100 kb apart. Genome maps without CGR but whose haplomap partner harbor at least one CGR were also included.

Alignments of fusion maps to hg38 identified reference genomic regions contributing to the composition of the 101 genome maps. To find the most represented reference fragment in the fusion maps, adjacent aligned regions were first merged (resulting in 100 reference donor fragments), and the total number of fusion map alignments summed for each reference donor fragment, excluding duplicate counting from haplomap pairs. Two reference regions, Chr1:188,188,529-189,139,998 and Chr12: 68,713,897-69,940,974, were found in at least 10 pairs of fusion haplomaps. The sum of fusion map alignments across the whole genome was used as the copy number profile for the 101 fusion maps in the circos plot (Fig. 7).

Piecewise alignments of optical maps to distinct and distant reference genomic regions are a signature of complex genomic rearrangements. A genomic fusion is characterized by a pair of neighboring alignments, on an optical map, to distinct regions of the reference genome, with breakpoints corresponding to the closest pair of aligned labels. Often, a fusion junction contains a stretch of DNA (label patterns) that does not align anywhere on the reference genome. These unknown regions likely correspond to significantly rearranged reference fragments (potentially with additional fusions) that no longer bear semblance to the original reference (examples in Fig. 6B and Supplementary Fig. 7).

Genomic distributions of translocations and large insertions and deletions on the 101 fusion maps were calculated as the total number of each SV type per 1 Mb windows (Supplementary Fig. S5). This analysis excluded all SV overlapping N-base gaps and segmental duplications on hg38.

### Statistical Analyses

The Wilcoxon rank sum test with continuity correction was used to assess the null hypothesis that insertion and deletion sizes have the same distributions (Fig. 3B). The null was significantly rejected for both all genome maps (p<10^-10^) and for the 101 fusion maps (p=0.005; Supplementary Fig. S4). The Welch two-sample t-test was used to assess the null hypothesis that fusion junction intervals (distance between a breakpoint pair on the fusion map) for fusions found in the fusion maps there were and were not also identified from deep sequencing of neochromosomes (Supplementary Fig. S9). The null hypothesis was rejected at P<10^-7^.

The two-sample Kolmogorov-Smirnov test was used to evaluate whether neighboring fusion breakpoint distances on the reference genome deviated from the null hypothesis of random distribution (Korbel and Campbell 2013) (Fig. 8).

### Use of neochromosome deep sequencing data

Two supplementary data files from Garsed et al. 2014 were used in this study: Table S2 (worksheet “778 (DR)”) containing a list of genomic fusions and Table S3 (worksheet “778_CN”) containing the copy number profiles of the two neochromosome isoforms of cell line 778. As the original study was performed with hg19, genomic coordinates in these data were remapped to hg38, using the NCBI Genome Remapping Service (https://www.ncbi.nlm.nih.gov/genome/tools/remap). Remappings to non-assembled chromosomes (chromosomes designated with suffixes _random and _alt) and unplaced sequences (chrU) were excluded. This resulted in the unsuccessful remapping of: (i) two fusions because one of their breakpoint pairs maps ubiquitously to three different assembled chromosomes; (ii) one fusion because one of its breakpoints cannot be remapped to hg38, and (iii) one CGR because it cannot be remapped to hg38.

The copy number profile and genomic fusions in the circos plot of Figure 7 was generated based on the remappings of these two tables.

A fusion on a fusion map is considered as concordant with a fusion on the neochromosomes (per Table S2: 778 (DR) of Garsed *et al.* 2014) if both breakpoints of the genome map fusion are within 100 kb of the neochromosome fusion breakpoint pair.

## Data Access

The Bionano Solve output for cell line 778 is available in the Supplementary Information.

## Acknowledgements

We thank Arcadi Cipponi from Garvan Institute of Medical Research for preparing the cell line and Joyce Lee and Alex Hastie from Bionano Genomics Inc., San Diego, USA, for providing update analysis with Bionano Solve version 3, prior to official release. This work was funded by the by Movember Australia and the Prostate Cancer Foundation Australia as part of the Movember Revolutionary Team Award to the Garvan Institute of Medical Research. V.M.H. is supported by the University of Sydney Foundation and Petre Foundation, Australia.

## Disclosure Declaration

The authors declare no competing financial interest.

